# A Conserved Domain of *Cfap298* Governs Left-Right Symmetry Breaking in Vertebrates

**DOI:** 10.1101/2025.05.16.654593

**Authors:** Marvin Cortez, Cullen B Young, Katherine A Little, Daniel T Grimes, Danelle Devenport, Rebecca D Burdine

## Abstract

*Cfap298* is a highly conserved gene required for ciliary motility and dynein arm assembly. It plays a known role in Left-Right (LR) patterning in zebrafish and is linked to human ciliopathies. Here we describe a new *Cfap298* mutant allele, *Cfap298^Δ3aa^*, which selectively interferes with LR axis establishment in mice. Mutant embryos display a range of laterality defects including *situs solitus*, *situs inversus*, and *situs ambiguous,* as indicated by abnormal heart, lung, and stomach positioning. At embryonic day 8.5, mutant embryos display abnormal *Nodal*, *Pitx2*, and *Lefty1* expression patterns in the lateral plate mesoderm and midline, consistent with an early disruption in LR symmetry breaking. In mice, LR asymmetry is established by leftward fluid flow in the node, generated by planar-polarized cilia. Although *cfap298* mutations are reported to affect cell polarity, we did not observe changes in cilia position, length, or planar cell polarity protein localization within the node, suggesting that *Cfap298^Δ3aa^*functions at the level of cilia motility. Consistently, motile cilia lining the trachea of *Cfap298^Δ3aa^* mutants fail to beat. Expression of the *Cfap298^Δ3aa^* variant in zebrafish fails to rescue the LR defects of *cfap298* (*kurly)* loss-of-function mutants. These results confirm that *Cfap298* functions in LR axis formation in mammals and uncover a novel region of CFAP298 protein with a conserved and essential role in cilia motility.

**Summary Statement:** A novel *Cfap298* mutation disrupts left-right patterning in mice by impairing ciliary motility, revealing a conserved protein region essential for this function across species.

## Introduction

Motile cilia are microtubule-based structures used by many organisms throughout the animal kingdom to move fluids or generate flow. In mammals, motile cilia line the trachea to clear the airways while motile cilia on ependymal cells propel cerebral spinal fluid throughout brain ventricles. Motile cilia in the oviduct generate flow to move oocytes towards the uterus and form the flagella that propel sperm towards the oocyte (Spassky and Meunier, 2017; Hilgendorf et al., 2024). Dysregulation of motile cilia function in these tissues can lead to cystic fibrosis, hydrocephalus, and infertility in men and women. These disorders are part of a larger class of congenital disorders known as ciliopathies (Spassky and Meunier, 2017; Hilgendor et al., 2024).

Motile cilia also play critical roles in establishing the Left-Right (LR) body axis during development in humans, mice, *Xenopus* and zebrafish. Motile cilia in structures collectively known as Left-Right Organizers (LROs) generate fluid flows that culminate in the asymmetric expression of the TGFβ ligand Nodal on the left in the lateral plate mesoderm (LPM) (Grimes and Burdine, 2017; Blum and Ott, 2019; Little and Norris, 2021; Forrest et al., 2022). The mouse LRO is the node, a transient pit-like embryonic structure with planar polarized motile cilia that move fluid across the node to the left side. Crown cells surrounding the node are proposed to sense the leftward fluid flow using immotile cilia (Grimes and Burdine, 2017; Hashimoto et al. 2010). In zebrafish, the LRO is Kupffer’s vesicle, a transient fluid-filled epithelial sac where motile cilia generate a counterclockwise flow. Mutations in cilia motility genes perturb fluid flow in LROs disrupting left-sided *Nodal* expression, resulting in abnormal organ positioning within the body cavity known as *situs inversus* (mirror image arrangement) or heterotaxy (random arrangements) (Grimes and Burdine, 2017).

One gene that has been previously shown to be essential in cilia motility is the cilia and flagella-associated protein 298 (CFAP298) (Austin et al., 2013; Jaffe et al., 2016). CFAP298 is required in the cytoplasm for proper preassembly of axonemal dyneins needed for cilia motility and thus, LR patterning (Wang et al., 2022; Jaffe et al., 2016; Austin-Tse et al., 2013). Mutations in the zebrafish ortholog *kurly* lead to loss of inner and outer dynein arms within the ciliary axoneme resulting in immotile cilia and LR patterning defects (Jaffe et al., 2016). Furthermore, mutations in human CFAP298 have been identified in patients with heterotaxia (Austin-Tse et al., 2013). Work further suggests CFAP298 may play a role in planar cell polarity (PCP) based on cilia polarity in the zebrafish kidney and loss of asymmetric PCP protein localization in *Xenopus* larval skin cells (Jaffe et al., 2016), but this role has not been further elucidated. Utilizing CRISPR-Cas9 we generated a mutation that alters three amino acids in a highly conserved region in the mouse ortholog of *Cfap298, Cfap298^Δ3aa^. Cfap298^Δ3aa^* embryos display *situs inversus* and heterotaxia accompanied by abnormal expression of *Nodal*, *Pitx2*, and *Lefty1*. We show *Cfap298^Δ3aa^* specifically affects cilia motility, and not planar cell polarity, in the mouse node. Rescue experiments in zebrafish demonstrate that while *Cfap298^Δ3aa^* mRNA retains some function, it is unable to rescue LR defects in *cfap298* mutant zebrafish embryos. Thus, the *Cfap298^Δ3aa^* allele identifies a three amino acid region essential for its cilia motility related functions.

## Results

### *Cfap298^Δ3aa^* mutants display defects in organ laterality

*Cfap298* (*C21ORF59*; *kurly; FBB18*) encodes a small, 290 amino acid protein containing a coiled-coil domain (CCD) and an uncharacterized domain of unknown function (DUF) (Austin et al., 2013). Recently, a ubiquitin-like (UBL) domain and Loop domain were characterized in the *Chlamydomonas* ortholog *FBB18* (Yamamoto et al., 2025). To investigate the functional role of *Cfap298* in mouse development, we isolated a novel allele, *Cfap298^Δ3aa^*, using CRISPR-Cas9 genome editing. The allele contains a six-nucleotide deletion together with two nucleotide substitutions in exon 4, resulting in the deletion of two amino acids (Y161, D162) and a missense mutation (P163S) at an adjacent residue (Fig. 1A). The three affected amino acids map to an alpha helix adjacent to the Loop domain and just before the UBL domain (Fig. 1B, Fig S1B). Sequence conservation analysis using DMfold (deepMSA2; Zheng et al., 2024) identified residues D162 and P163 as evolutionarily conserved (Fig. S1A).

**Figure 1.**
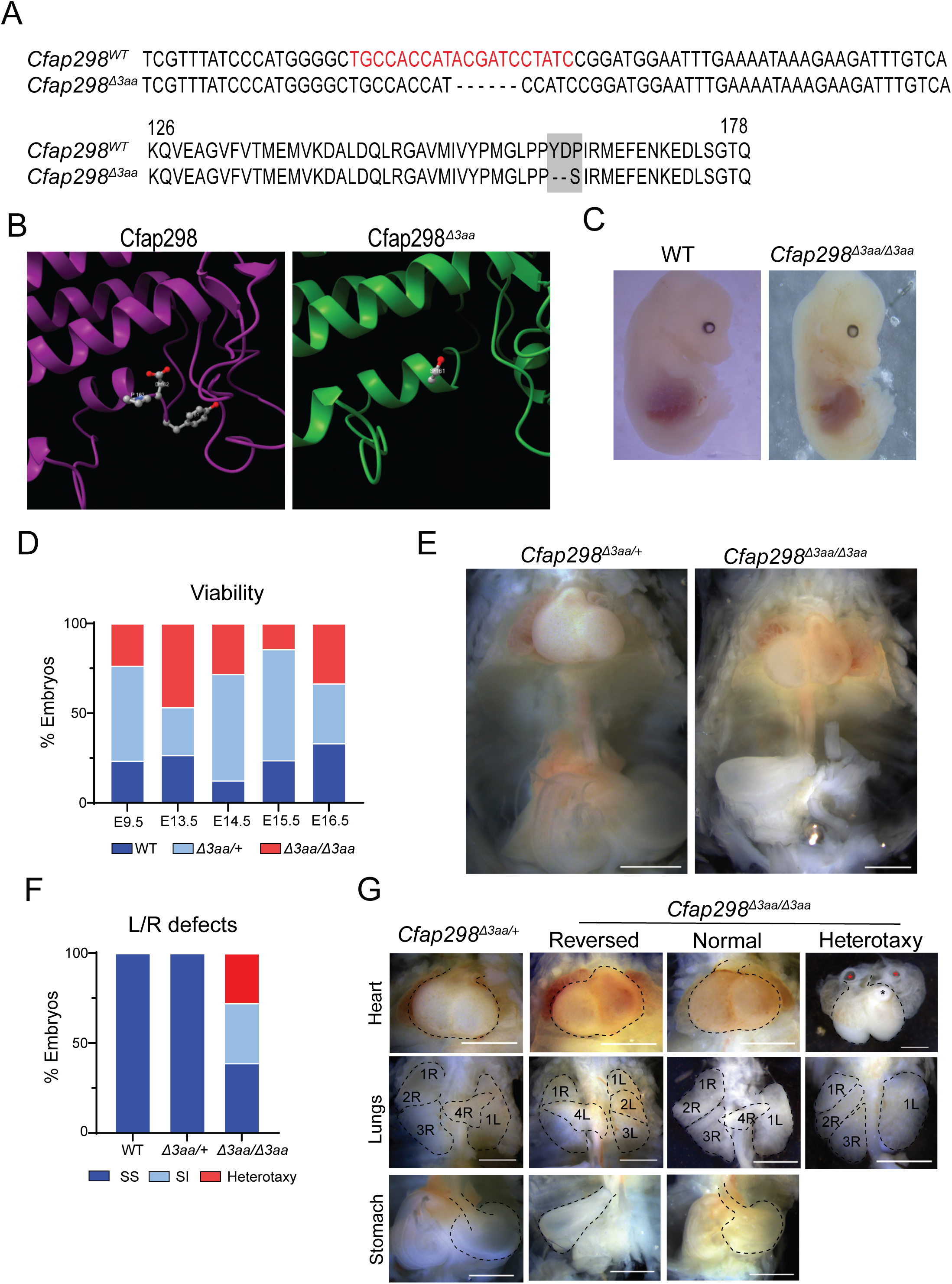
*Cfap298^Δ3aa^*mutants display Left-Right laterality defects. A) Generation of the *Cfap298^Δ3aa^*mutant allele by targeting exon 4 of the *Cfap298* (*Cfap298*) gene using CRISPR/Cas9 gene targeting. Guide recognition sequence within the exon 4 sequence is highlighted in red on the wildtype sequence. The *Cfap298^Δ3aa^*mutation consists of a 6 base deletion and a base substitution of T->C, resulting in removal of 2 amino acids at positions 161-162 and a missense mutation of P163S. B) AlphaFold predicted structures of Cfap298 and Cfap298^Δ3aa^ proteins in the region of the Δ3A mutation. Y161 and D162 are labeled on wildtype Cfap298, and S161 is labeled on Cfap298^Δ3aa^. C) Whole embryo images of E14.5 wildtype and *Cfap298^Δ3aa^*^/*Δ3aa*^ embryos. Scale bar: 1mm. D) Viability distribution of wildtype, heterozygous, and homozygous mutant embryos at E9.5 (n=17), E13.5 (n=15), E14.5 (n=33), E15.5 (n=21) and E16.5 (n=6). E) Representative images of body cavities showing heart, lungs and stomach positions from *Cfap298^Δ3aa^*^/+^ and *Cfap298^Δ3aa^*^/*Δ3aa*^ E14.5 embryos. F) Quantification of *situs solitus*, *situs inversus*, and heterotaxy from wildtype (n=21), *Cfap298^Δ3aa^*/+ (n=46), and *Cfap298^Δ3aa^*^/*Δ3aa*^ (n=18) embryos from E13.5-E15.5 stages. G) Representative images of heart, lung and stomach from E14.5 *Cfap298^Δ3aa^*^/+^ and *Cfap298^Δ3aa^*^/*Δ3aa*^ embryos displaying with *situs solitus* (Normal), *situs inversus* (Reversed) and heterotaxy phenotypes. Hearts and stomachs are outlined. Individual lung lobes are outlined and labeled with their position on either left (L) or right (R). Atriums (red asterisk) and ventricle (black asterisk) are indicated in heart with heterotaxy. Scale bar: 1mm.

To examine developmental consequences of this mutation, we analyzed embryos at embryonic day 14.5 (E14.5). Homozygous mutants (*Cfap298^Δ3aa/Δ3aa^*) displayed no overt morphological abnormalities externally and were recovered at expected Mendelian ratios, indicating that embryonic viability at mid-gestation was unaffected (Fig. 1C, D). Given the established role of *Cfap298* orthologs in left-right (LR) symmetry breaking (Austin-Tse et al., 2013; Jaffe et al., 2016), we assessed internal organ situs from E13.5 to E15.5. Whereas wild-type (WT) and *Cfap298^Δ3aa/+^* heterozygous embryos exhibited exclusively *situs solitus*, homozygous mutants displayed a significant incidence of LR patterning defects (60%, 11/18 embryos), including complete *situs inversus* (Fig. 1E-G; Table 1). Despite the high incidence of LR defects, we were able to raise one *Cfap298^Δ3aa^*mutant to adulthood. This mutant displayed heterotaxy including a reversed heart, right mirrored liver, left sided stomach, and left lung isomerism compared to a *Cfap298^Δ3aa^*^/+^ adult which displayed normal situs of these organs (Fig. S2B). These findings align closely with the previously reported phenotypes of zebrafish *kurly* mutants, which similarly display randomized visceral organ placement due to impaired motile cilia function in Kupffer’s vesicle (Jaffe et al., 2016). Consistent with LR abnormalities, we observed congenital heart defects (CHDs) in *Cfap298^Δ3aa^* homozygous mutant embryos, including Transposition of the Great Arteries (TGA), Double Outlet Right Ventricle (DORV), and pulmonary stenosis (Fig. 1G).

**Table 1.** Stomach, heart, and lung situs *Cfap298^Δ3aa^* mutant embryos. Representative multi-organ situs observed in embryos. *Situs solitus, situs inversus,* and *heterotaxy* were determined by characterizing stomach, heart, and lung situs. Normal stomach situs was determined as being oriented towards the left side of the body and reversed as determined as oriented towards the right side. Normal heart situs was determined by orientation of the heart apex towards the left side of the body and reversed as oriented to the ride side. Normal lung situs was determined as the left side of the lung having 1 lobe, and the right side having 4 lobes and reversed as have 4 lobes on the left side and 1 lobe on the right. In one instance, a heart was observed to be left sided but also observed to have TGA. In one instance, an embryo had an abnormal lung having 3 lobes on the right side and one lobe on the left.

| *Embryo#* | *Genotype* | *Stomach situs* | *Heart situs* | *Lung situs* |
| --- | --- | --- | --- | --- |
| 1 | MUT | Normal | Reversed | abnormal |
| 2 | MUT | Normal | L-TGA | normal |
| 4 | MUT | Reversed | Reversed | reverse |
| 5 | MUT | Normal | Normal | normal |
| 5 | MUT | Reversed | Reversed | Reversed |
| 8 | MUT | Reverse | Reversed | Reversed |

### *Cfap298^Δ3aa^* mutants fail to establish correct expression of LR patterning genes

The appearance of LR defects in organ laterality is indicative of perturbed LR patterning earlier in embryogenesis. Thus, we next assessed the asymmetric expression of key genes involved in executing the LR laterality program including *Nodal* and its target *Pitx2* in the lateral plate mesoderm (LPM) of E8.5 embryos. Spatial expression analysis by Hybridization Chain Reaction (HCR) using probes against *Nodal, Pitx2*, and *Lefty1* revealed left-sided *Nodal*, *Pitx2* and *Lefty1* in the LPM and *Lefty1* expression in the midline in wildtype embryos, as expected (Fig. 2A). By contrast, *Cfap298^Δ3aa^*homozygous mutant embryos exhibited disrupted *Nodal*, *Pitx2*, and *Lefty1* expression (Fig. 2B). In some embryos, we observed right-sided expression of *Nodal* and *Pitx2* in the LPM (Fig. 2B’). In another embryo, we observed weakened bilateral *Nodal* together with right-sided expression of *Pitx2* in the LPM and a loss of *Lefty1* at the midline (Fig. 2B’’). Further, in some embryos, *Nodal* and *Pitx2* expression was lacking in the LPM, but expression of *Lefty1* at the midline was normal (Fig. 2B’’). We conclude from these data that the LR organ laterality defects observed in E14.5 *Cfap298^Δ3aa^* mutant embryos are a result of the failure to correctly establish the LR axis.

**Figure 2.**
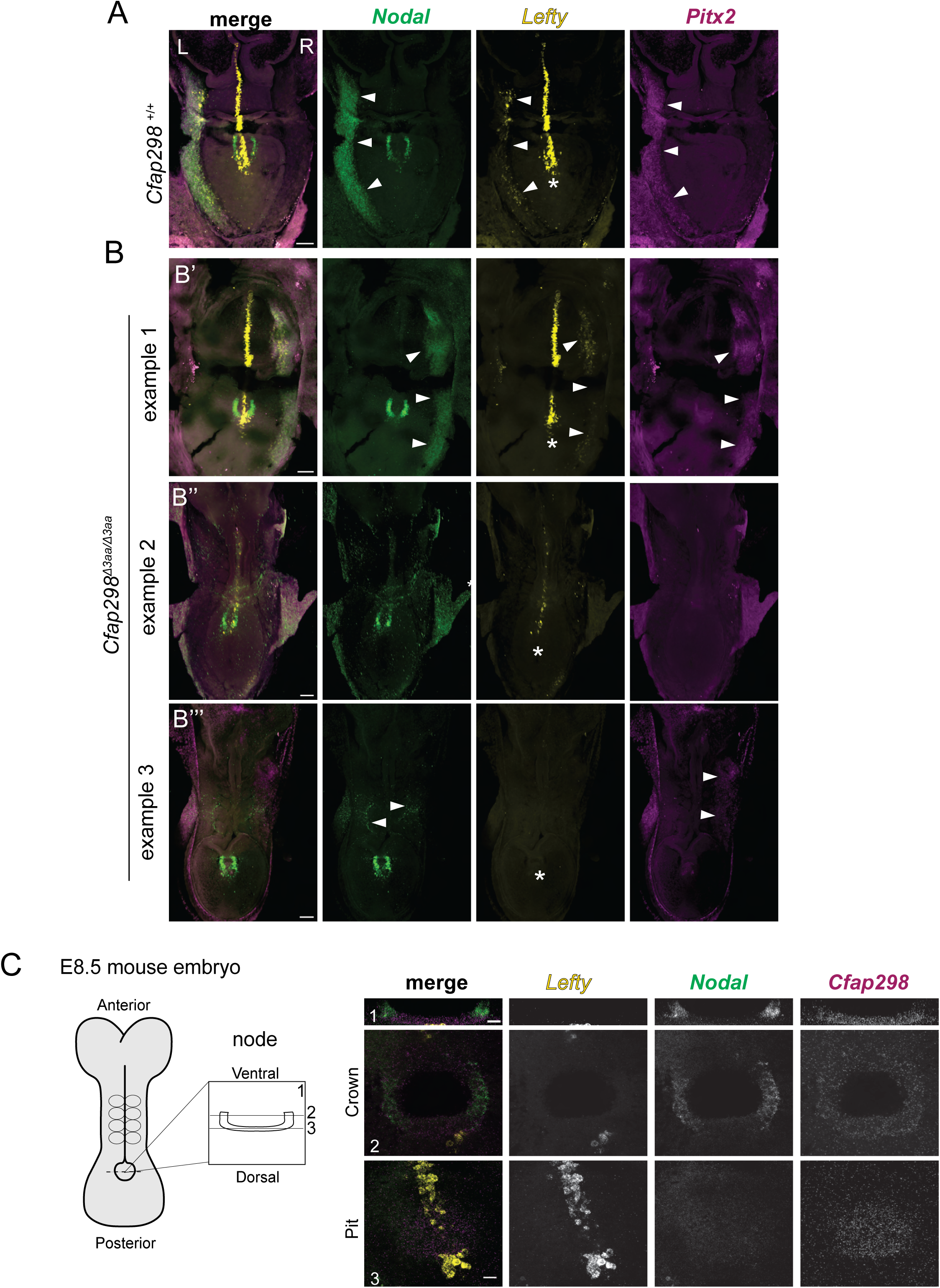
*Cfap298^Δ3aa^*mutants have perturbed Left-Right patterning. A) Representative HCR images for *Nodal* (green), *Pitx2* (magenta), and *Lefty1* (yellow) expression in whole-mount E8.5 wildtype embryo (A) and *Cfap298^Δ3aa^*mutants (B’, B’’, B’’’). Wildtype embryos exhibit left-sided *Nodal* and *Pitx2* expression in the lateral plate mesoderm (LPM) (arrowheads) with *Lefty1* expression along the embryonic midline (*). B) Images from *Cfap298^Δ3aa^*mutants showing one example of right-sided *Nodal, Lefty* and *Pitx2* expression along the LPM (arrowheads) (B’), an example where *Nodal* and *Pitx2* are absent in the LPM and *Lefty1* is expressed normally at the midline (B’’), and an example of abnormal expression with reduced and bilateral *Nodal*, right-sided *Pitx2* (arrowheads), and absent *Lefty1* in the midline (*) (B’’’). Scale bar: 100um. C) Diagram of E8.5 wildtype embryo viewed from the dorsal side, anterior is up. Expanded view of a cross section through the node is shown on the right. Representative images of E8.5 node labeled by HCR for *Nodal* (green) *Lefty1* (yellow) and *Cfap298*(magenta) transcripts. Cross section (1) and planar (2,3) views of the node showing *Nodal* expression in the crown cells but not at the pit cells. *Lefty1* expressing cells mark the floorplate of the neural tube are visible at the base of the pit of the node. *Cfap298* expressing cells are visible throughout the node. Scale bar: 10um.

### Node morphogenesis and cilia assembly are unaffected in *Cfap298^Δ3aa^*mutants

LR patterning requires an intact LRO, motile cilia, and proper planar polarity to align cilia. Single-cell RNA sequencing (scRNA-seq) datasets from E8.5 mouse embryos (Pijuan-Sala et al., 2019) revealed that while *Cfap298* transcripts are ubiquitously expressed, they are highly enriched within the node/notochord cell population, similar to the LR patterning gene, *Nodal*, and consistent with expression patterns previously observed for zebrafish *kurly* (Jaffe et al., 2016; Fig. S3). Using HCR, we confirmed the widespread expression of *Cfap298* transcripts as well as elevated levels specifically within crown and pit cells of the embryonic node at E8.5 (Fig. 2C). The conserved expression pattern of *Cfap298* in the node suggests an essential and early role in LR symmetry breaking.

To determine the cause of LR patterning defects in *Cfap298^Δ3aa^*mutants, we investigated the formation of the node and cilia. The node is a transient, pit-like structure located at the posterior midline of E8.0-E8.5 embryos. It is lined with motile cilia that generate a leftward fluid flow, initiating the symmetry-breaking event in establishment of the LR axis (Hamada, 2020). At E8.5, morphologically normal nodes were present in *Cfap298^Δ3aa^* mutant embryos, and their overall shape were similar to wildtype littermates (Fig. 3A). To determine whether cilia assembly was altered in *Cfap298^Δ3aa^* mutants, we measured the length of nodal cilia labeled with antibodies against acetylated tubulin but did not observe an appreciable difference in cilia length between wildtype and mutant embryos (Fig. 3B).

**Figure 3:**
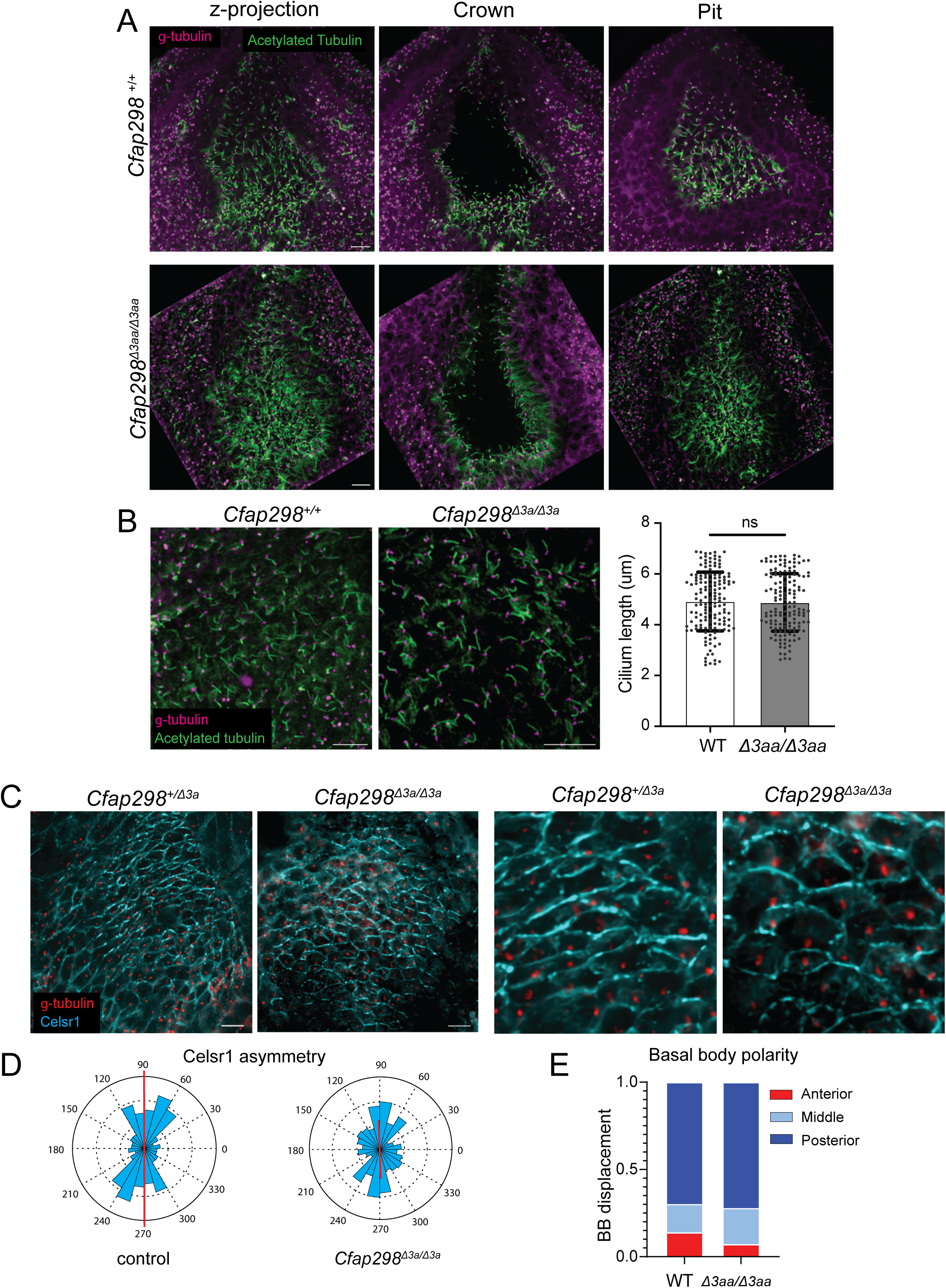
*Cfap298^Δ3aa^* mutants develop normal nodes. A) Representative immunofluorescent images of E8.5 nodes from wildtype and *Cfap298^Δ3aa^*mutant embryos. Embryos are stained for cilia with acetylated tubulin (green) and gamma tubulin (magenta). For wildtype and mutant nodes, one plane is shown for the crown cells and another plane shown for pit cells along with a z-projection of the whole node. B) Images of cilia are shown for wildtype and mutant nodes. Quantifications of cilium length for wildtype (n=3 nodes, 150 cilia total) and mutant (n=3 nodes, 150 cilia total) embryos showing no significant difference (p=0.76 by unpaired t test). C) Images showing Celsr1 (cyan) and gamma-tubulin (red) marking basal bodies in WT and *Cfap298^Δ3aa^*mutant embryos. Circular histograms display magnitude and orientation of Celsr1 polarity along the AP axis of wildtype (n=3 nodes, 99 cells total) and mutant (n=3, 218 cells total) embryos. D) Quantification of basal body polarity along the AP axis of the wildtype (n=3, 86 cells total) and mutant (n=3, 97 cells total) embryos. Scale bars=10um. ns= not significant.

### *Cfap298^Δ3aa^* does not disrupt PCP establishment

In multiciliated cells of *Xenopus* epidermis, *Cfap298* mutations disrupt the asymmetric localization of the core PCP component, Prickle (Pk) (Jaffe et al., 2016), suggesting that *Cfap298* may function in PCP establishment. To determine if the *Cfap298^Δ3aa^* mutation affects PCP, we first examined PCP establishment in the developing epidermis, a well-established system to investigate PCP function in mouse embryos (Devenport and Fuchs, 2008; Aw et al, 2016). Although mouse skin does not contain cells with motile cilia, *Cfap298* is broadly expressed in most cell types of the skin at E14.5 including epidermal cells, hair placodes, dermal fibroblasts and the dermal condensate (Figure S4A) (Sennet el al., 2015; Rezza et al., 2016). Thus, the skin provides an ideal system to evaluate the role of *Cfap298* in PCP without the confounding effects of cilia-driven fluid flow, which can orient and align planar polarity through positive feedback (Mitchell et al, 2007; Guirao et al., 2010). We therefore measured the asymmetric localization of the core PCP protein Celsr1 in basal cells of the epidermis, as well as the polarization and alignment of hair follicles along the anterior-posterior (AP) axis, which is the downstream output of PCP asymmetry in mammalian skin (Devenport and Fuchs, 2008). However, we did not observe a reduction in the asymmetry of Celsr1, which was enriched along the AP junctions of basal epidermal cells in both control and *Cfap298^Δ3aa^*mutant embryos (Fig. S4B). Moreover, we did not observe defects in hair follicle polarity or alignment (Fig. S4C), indicating that PCP establishment occurs normally in the epidermis of *Cfap298^Δ3aa^* mutants. Since *Cfap298* has been implicated in planar polarization of motile cilia in other organisms (Jaffe et al., 2016), we investigated the impact of the *Cfap298^Δ3aa^*mutation on planar cell polarity (PCP) establishment and polarized basal body positioning within the node. Components of the PCP pathway, including Vangl1/2, Dvl2/3, Pk2 and Celsr1, are asymmetrically localized along the AP axis of the node where they govern the posterior displacement of the basal body, important for leftward fluid flow (Hashimoto et al., 2010, Antic et al., 2010, Mahaffey et al., 2016, Minegishi et al., 2017). Defects in PCP within the node affect cilia placement and fluid flow leading to defects in LR patterning. To determine whether PCP is established correctly in the nodes of *Cfap298^Δ3aa^* mutants, we measured the orientation and magnitude of Celsr1 asymmetry (Aw et al., 2016). In wildtype embryos, Celsr1 was enriched along the AP junctions of node epithelial cells, as expected. In *Cfap298^Δ3aa^* mutant embryos, Celsr1 was similarly asymmetrically polarized along the AP junctions (Fig. 3C). Downstream of PCP establishment in the node, basal bodies become asymmetrically positioned toward the posterior of each cell (Hashimoto et al., 2010). We therefore measured the relative distance of basal bodies, marked by gamma-tubulin, along each cell’s AP axis. In wildtype embryos, basal bodies were positioned toward the posterior end of pit cells as expected. In *Cfap298^Δ3aa^*mutants, basal bodies were also posteriorly polarized (Fig. 3D). Collectively, these observations indicate that node morphogenesis and planar polarization occur normally in *Cfap298^Δ3aa^* mutants, and that LR defects in mutant embryos must occur downstream or independent of cilia axoneme assembly or planar polarity.

### *Cfap298^Δ3aa^* mutants have immotile cilia

Given the lack of PCP defects in the node, we hypothesized that the Δ3aa mutation may be impacting cilia motility function in mice. To test this, we imaged the trachea, which is lined with multiciliated cells (MCCs) in a control and a mutant adult mouse. The adult mutant seemed to display hydrocephalus indicative of impaired cilia motility in the brain (Spassky and Meunier, 2017; Fig. S2A). We observed MCCs in both *Cfap298^Δ3aa/+^* and *Cfap298^Δ3aa^*mutant tracheas (Movies 1 and 2). However, while MCCs in *Cfap298^Δ3aa^*^/+^ mice were highly motile, we observed severely reduced to no movement of cilia in the mutant trachea (Movies 1 and 2). This shows that the *Cfap298^Δ3aa^*mutation disrupts cilia motility.

### *Cfap298^Δ3aa^* does not rescue LR defects in zebrafish

Our results thus far indicate that amino acids Y161, D162 and P163 lie within a region of the mouse CFAP298 protein that functions in LR patterning and cilia motility. To determine if these residues perform a conserved function in LR asymmetry in vertebrates, we tested whether the *Cfap298^Δ3aa^* variant was capable of rescuing zebrafish *cfap298* null mutants. The *cfap298^tj271^*allele was previously shown to be a loss-of-function mutation, and zebrafish embryos homozygous for *cfap298^tj271^* (hereafter referred to as *cfap298*^−/−^) display several phenotypes associated with cilia motility defects including body and tail curvature, randomized heart jogging and kidney cysts (Jaffe et al., 2016) (Fig. 4A, C). Injection of *cfap298*^−/−^ embryos with 500pg of wildtype zebrafish *cfap298* mRNA fully rescued both body curvature and LR defects, as determined by scoring heart jogging at 48hpf (Fig. 4C, D). By contrast, injection of *cfap298*^−/−^ embryos with zebrafish *cfap298^Δ3aa^*mRNA, which contains the orthologous *Δ3aa* amino acid changes (ΔH161, ΔD162, and P163S), failed to rescue LR defects (Fig. 4B, C). Interestingly, 40% (11/27) of *cfap298^−/^*^−^ embryos injected with *cfap298^Δ3aa^*mRNA displayed normal body curvature (Fig. 4C, D). Although the cellular basis of body curvature phenotypes in zebrafish cilia mutants is not understood (Bearce and Grimes, 2021), it appears that *cfap298^Δ3aa^* can partially execute its function in maintaining body and tail straightness suggesting *Cfap298^Δ3aa^* is a partial loss-of-function mutation. Taken together our evidence suggests that the mutation in *Cfap298^Δ3aa^* defines a three amino-acid region critical for cilia motility.

**Figure 4:**
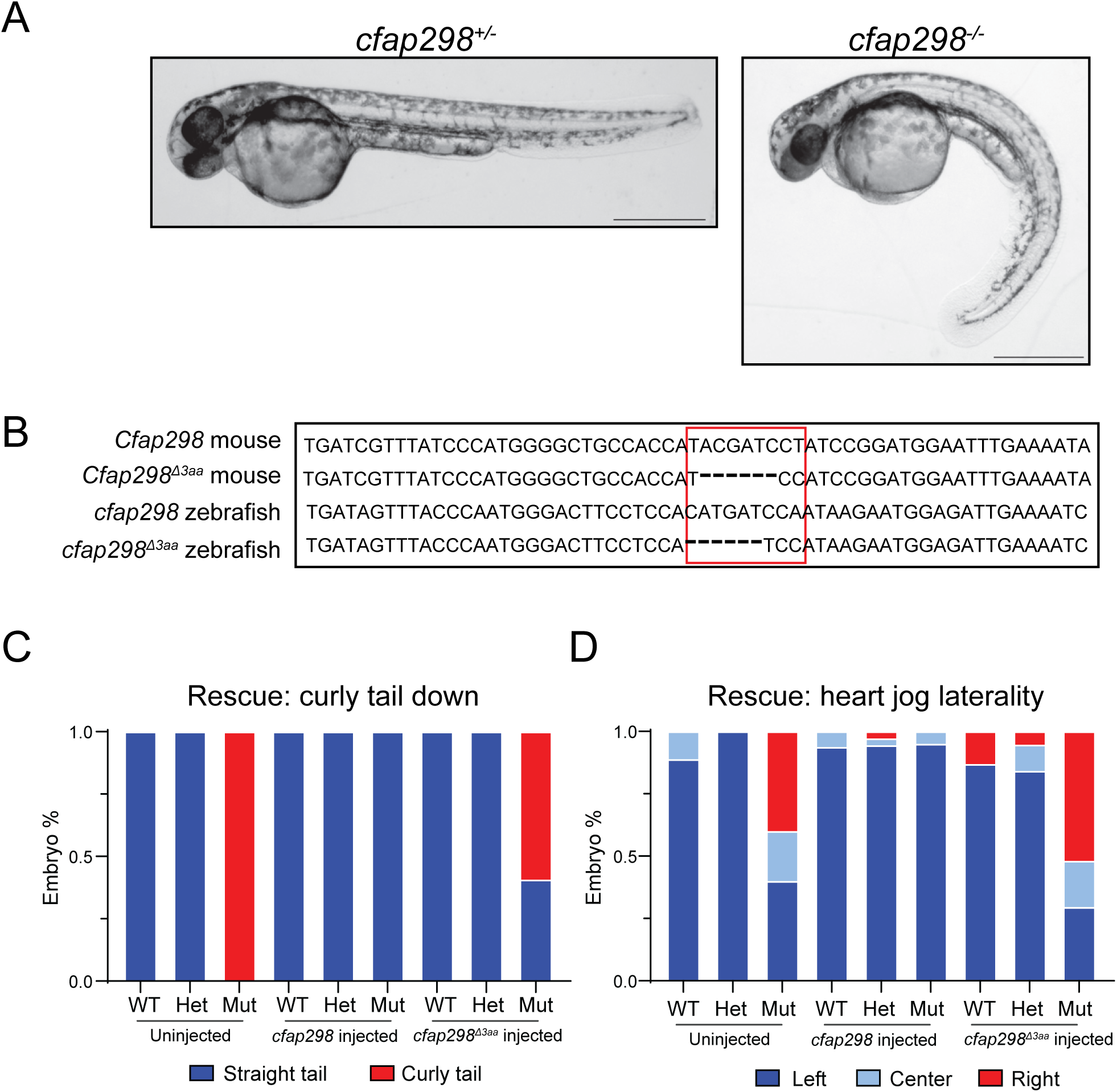
*Cfap298^Δ3aa^*does not rescue LR defects in zebrafish. A) Representative images of *cfap298*^+/−^ and *cfap298*^−/−^ zebrafish embryos at 48hpf, with *cfap298*^+/−^ embryos displaying straight bodies/tails compared to *cfap298*^−/−^ which displaying curved bodies/tails. B) Comparison of mouse wildtype *Cfap298* and *Cfap298 ^Δ3aa^*with wildtype zebrafish *cfap298* and engineered *cfap298 ^Δ3aa^* sequence. C-D) Quantification of body curvature (C) and heart position (D) observed in uninjected and injected embryos. Uninjected wildtype (n=9) and *cfap298*^+/−^ (n=13) embryos were observed to have straight tails and few or no heart jogging defect compared to *cfap298*^−/−^ (n=10) which all displayed curly tails and increased heart jogging defects. Similarly, wildtype (n=16) and *cfap298*^+/−^(n=36) embryos injected with 500pg of wildtype *cfap298* mRNA were observed to have straight tails and no heart jogging defects, however, *cfap298*^−/−^ (n=20) embryos were observed to have straight tails along with normal heart jogging. Wildtype (n=23) and *cfap298*^+/−^(n=38) embryos inject with 500pg of *cfap298^Δ3aa^* mRNA displayed straight tails and few heart jogging defects. *cfap298*^−/−^ (n=27) embryos injected with *cfap298^Δ3aa^* mRNA were observed to have a partial rescue in body curvature with some mutants showing straight tail phenotypes but were not observed to have rescued heart jogging defects. Scale bar 500um.

## Discussion

The mechanisms underlying LR symmetry breaking are highly conserved across vertebrates, and the involvement of motile cilia in the process is well established. In previous work, we demonstrated an essential role for *Cfap298* in LR symmetry breaking in zebrafish through its function in cilia motility (Austin et al., 2013; Jaffe et al, 2016). Here, through the use of a novel mutant allele, *Cfap298^Δ3aa^*, we reveal that *Cfap298* is also essential in LR axis formation in mice. Remarkably, changes in just three amino acids of the CFAP298 protein result in severe defects in LR organ positioning, preceded by abnormal LR patterning as determined by *Nodal*, *Pitx2*, and *Lefty1* expression in the LPM and midline. Given that ciliogenesis and planar polarization within the LRO are largely unaffected in *Cfap298^Δ3aa^*mutant embryos, we conclude that defects in LR symmetry breaking are due to a loss of cilia motility and fluid flow. Although we have not directly visualized cilia movement or fluid flow within the node, the immotility of cilia within the trachea of a *Cfap298^Δ3aa^* mutant is consistent with this interpretation. Further, the failure of the *Cfap298^Δ3aa^* variant to rescue LR patterning defects in zebrafish point to a conserved role for this region of the CFAP298 protein in cilia motility.

The *Cfap298^Δ3aa^* mutation affects a region of the protein that has not been previously implicated in *Cfap298’s* cilia motility function. The Chlamydomonas ortholog of *Cfap298*, *FBB18*, and the zebrafish ortholog, *kurly*, have been shown to interact with flagellar and motile cilia dynein arm components as well as dynein arm assembly factors (Austin et al., 2013; Jaffe et al., 2016; Wang et al., 2022). Human pathogenic mutagenic alleles, which have been previously reported to truncate the CFAP298 protein near the C-terminal end of the protein, were also shown to disrupt dynein arm assembly (Austin et al., 2013; Wang et al., 2022). FBB18 interacts with chaperone proteins, and it has been posited that FBB18 is a co-chaperone for dynein assembly factors (Wang et al., 2022). Therefore, loss of cilia motility in *Cfap298^Δ3aa^* mutants are likely the result of lost or reduced dynein arm assembly. We hypothesize that amino acids Y161, D162 and P163S in CFAP298 may constitute an interaction site for components of the dynein assembly machinery or its chaperones. Future experiments will focus on defining which CFAP298 protein-protein interactions are altered by the Δ3aa mutation.

When CFAP298 was initially annotated, it was described as consisting of two domains, a Coiled-Coil Domain and a Domain of Unknown Function (DUF) (Austin et al., 2013). Recently, a crystal structure of the Chlamydomonas ortholog, FBB18, was solved showing it consists of a Ubiquitin-like (UBL) domain and a so-called Loop domain (Yamato et al., 2025). In relation to these domains, the *Cfap298^Δ3aa^* mutation is predicted to affect a short alpha-helix located in the linker between Loop and UBL domains. How this region could be working in conjunction with the UBL and Loop domain of CFAP298 in dynein arm assembly also warrants future investigation.

Planar polarization of nodal cilia is important for directed cilia motility and the production of leftward fluid flow (Hashimoto et al., 2010; Song et al., 2010) and several pieces of evidence had previously implicated CFAP298 in PCP establishment. Zebrafish Kurly interacts with the core PCP protein, Dvl2, and kidney cilia lose planar polarization in *kurly* mutants (Jaffe et al., 2016). Further, depletion of CFAP298 disrupts the asymmetric localization of Prickle in multicilated cells in Xenopus, suggesting a potential upstream role for CFAP298 in PCP establishment (Jaffe et al., 2016). However, fluid flow itself can serve as an orienting cue for PCP and cilia motility is known to positively feedback on planar polarity (Mitchell et al, 2007; Guirao et al., 2010). Thus, the PCP defects observed in tissues with motile cilia could be a secondary consequence of defective cilia motility. We therefore examined PCP establishment in tissues with and without motile cilia to determine whether the effects of *Cfap298^Δ3aa^* could be attributed to a loss of PCP. In the mouse epidermis, which is strongly planar polarized but lacks motile cilia, we did not observe defects in PCP establishment in *Cfap298^Δ3aa^* mutants, nor did we observe changes in PCP asymmetry or planar polarized basal body positioning in cells with motile cilia (the node). This supports our conclusion that the Δ3aa mutation selectively impairs its cilia motility-related function. It is possible that other regions of the CFAP298 protein are important for PCP establishment in mice. Further investigations using a loss-of-function allele would be required to determine the function of *Cfap298* in planar polarization.

It is interesting to note that *Cfap298* is broadly expressed in all cell types in mouse embryos as well as the skin, though most of these cell types lack motile cilia. Interestingly, loss of *Cfap298* was shown to sensitize cancer cell lines to DNA damage (Zhao et al., 2024). This suggests that *Cfap298* may have additional functions outside of cilia motility. If true, this would further support the conclusion that the Δ3aa mutation selectively impairs cilia motility without affecting other CFAP298 functions. Elucidating the function of *Cfap298* in cells without motile cilia is a promising direction for future studies.

## Acknowledgements

Philip Johnson and Laboratory Animal Resources staff for fish and mice care. Peter Romanienko in the Rutgers Cancer Institute Genome Editing Shared Resource Knockout mouse facility. Gary Laevsky and Sha Wang in the Princeton University Nikon Center of Excellence Confocal Core facility. Burdine and Devenport lab members for feedback.

## Methods and Materials

### Generation of the *Cfap298^Δ3aa^* mouse line, mouse husbandry and breeding

*Cfap298* deletion mutations were generated using CRISPR-Cas9. The CRISPR target sequence was TGCCACCATACGATCCTATC<u>CGG</u> (PAM site underlined) in exon 4. Tracr and crRNA (Sigma) were prepared with Cas9 protein (Sigma) and microinjected into C57BL/6J zygotes. Founders were determined to have insertions or deletions by Sanger sequencing and PCR (Genewiz). One founder was observed to have a 6bp deletion which removed 2 amino acids and changed Proline 163 to Serine. Founder was backcrossed to C57BL/6J mice once and then bred into a C3H/HeJ background. *Cfap298^Δ3aa^* carrying mice was outcrossed into the C3H/HeJ background a minimum of 10 generations and transmission of the mutation was determined by PCR. For PCRs, forward primer sequence was CCCAATGCACTTTCAGAAACA and reverse sequence was ACCCTGCCCCACATACCT. Genotyping PCR produces 2 bands for a heterozygous mouse with the *Cfap298^Δ3aa^*amplicon being 6 bases smaller than the wildtype. To generate *Cfap298^Δ3aa^*^/*Δ3aa*^ homozygous embryos, *Cfap298^Δ3aa^*^/+^ adult mice were mated. Staging of embryos was estimated based on the time a mating plug was observed (E0.5). *Cfap298^Δ3aa^* homozygous embryos were identified by a PCR showing only the mutant amplicon.

All animal procedures were approved by Princeton’s Institutional Animal Care and Use Committee (IACUC). All mice were housed in facilities accredited through the American Association for Accreditation of Laboratory Animal Care (AAALAC).

### E13.5-E15.5 embryo and adult necropsies

For embryo necropsies, E13.5-E15.5 embryos were dissected in 1XPBS. Tissue from tails were collected for genotyping. Whole embryos with incisions on the sides of the abdomen were then incubated in 4% PFA/PBS overnight at 4 degrees to allow for fixation of internal organs. Embryos were washed 2 times in PBS. For postnatal stages, pups were ear punched 12 days after birth and genotyped for *Cfap298^Δ3aa^*. For adult necropsies, mice were euthanized under CO2. Body cavities were opened to determine situs.

### E8.5 embryo Hybridization Chain Reaction (HCR)

The HCR procedure was adapted from Matthew Anderson and Mark Lewandoski, NIH, modified from Choi et al, Methods Mol Biol 2148, 159-178 (2020). E8.5 embryos were dissected in ice cold 1XPBS and then fixed overnight at 4 degrees in 4% PFA/PBS. Embryos were then washed 2 times for 5 minutes in PBS with 0.1% Tween20 (PBSTw), followed by a series of dehydration steps into 100% methanol. Embryos were incubated in methanol for at least 2 hours or overnight before proceeding with the probe step. Embryos were rehydrated into 100% PBSTw and then bleached in 6% Hydrogen peroxide in PBS for 20 minutes. Embryos were washed 2 times for 5 minutes in PBSTw while rocking. Embryos were then treated in a 1:1 mix of 10ug/ML Protenase K in PBSTw for 1 minute then washed 2 times for 5 minutes in PBSTw. The embryos were fixed again in 4%PFA for 20 minutes at room temp while rocking. Embryos were washed 3 times for 5 minutes in PBSTw, transitioned into 1:1 hybridization buffer warmed to 37 degrees and PBSTw for 10 minutes while rocking. Embryos washed for 10 minutes at room temperature in prewarmed hybridization buffer. Hybridization buffer was replaced with fresh buffer and embryos were incubated in hybridization chamber at 37 degrees while rocking for 1-3 hours. Embryos were then incubated in probe+hybridization buffer solution overnight in chamber. Embryos were washed in probe wash buffer warmed to 37 degrees 3 times for 20 minutes at 37 degrees while rocking. Embryos were rinsed in 5X Saline Sodium Citrate with 0.1% Tween20 (SSCT) 3 times for 5 minutes at room temperature while rocking. Embryos were then washed in 1:1 amplification buffer and PBSTw for 10 minutes at room temperature while rocking and then transferred into amplification buffer for 30 minutes at room temperature while rocking. Embryos were then incubated in hairpins prepared in amplification buffer overnight at room temperature away from light. One quick wash in 5xSSCT was followed by 3, 5 minutes washed in 5XSSCT at room temperature while rocking and away from light. Embryos were then whole mounted in SlowFade^TM^ Glass Soft-Set Antifade mounting media. After imaging, embryos were lysed in 95° C 50mM NaOH for genotyping.

Embryos were imaged on Nikon A1R-Si confocal microscopes. Images acquired were processed using NIS elements, ImageJ, and Adobe Photoshop.

### Immunofluorescence

For immunofluorescent labeling of the node, we followed an immunostaining protocol adapted from Sai et al. (2022). E8.5 embryos were dissected in cold PBS then fixed in 4% PFA/PBS on ice for 45 minutes. Embryos were washed 3 times for 10 minutes in PBS +0.2% Tritonx100 (PBST) then incubated in blocking solution (1% Bovine Serum Albumin and 2.5% Normal Donkey Serum prepared in PBST) for 1 hour at room temperature. After blocking, embryos were incubated in primary antibodies diluted in blocking solution overnight at 4 degrees. Following primary antibody incubation, embryos were washed 3 times for 15 minutes in PBST, followed by incubation in secondary antibodies and Hoescht prepared in PBST overnight at 4 degrees. Embryos were then washed 3 times for 5 minutes in PBS and whole mounted in SlowFade^TM^ Glass Soft-Set Antifade mounting media. After imaging, embryos were lysed in 95° C 50mM NaOH for genotyping.

For immunostaining of mouse embryonic skins, tails were first removed from E15.5 embryos for genotyping followed by fixation in 4% PFA/PBS with Mg^2+^ and Ca^2+^ for 1 hour at room temperature while rocking. Skins were then dissected from embryos and washed 3 times in PBST for 5 minutes each followed by incubation in blocking solution (PBST+ 1% fish gelatin, +1% Bovine Serum Albumin, 2.5% Normal Donkey Serum) for 1 hour at room temperature while rocking. Skins were then incubated overnight in primary antibody solution prepared in blocking solution overnight at 4 degrees while rocking. Skins were then washed 3 times in PBST for 30 minutes each. Washed skins were then incubated in secondary antibody and Hoescht solution prepared in PBST overnight at 4 degrees while rocking. Skins were washed 3 times for 10 minutes each followed by two 5-minute washes in PBS. Skins were mounted epidermal side up in Prolong Gold curing mounting media and imaged after curing.

Embryos were imaged on a Nikon A1R-Si confocal microscope. Images were processed using NIS elements, ImageJ, and Adobe Photoshop.

The following primary and secondary antibodies used: Celsr1 Guinea Pig 1:1000 (Devenport and Fuchs. 2008), acetylated-tubulin mouse 1:200 (Sigma Catalog# T6793), gamma-tubulin mouse 1:200 (Sigma catalog# T6557), P-cadherin rat (Invitrogen 13-2000z), Sox9 rabbit (Millipore AB5535), Alexa Fluor 488 Donkey anti-Guinea Pig (Jackson Immuno catalog# 706-545-148). Acetylated-tubulin primary antibody was directly conjugated with Alexa 555 fluorophore using Biotium Mix-n-Stain™ CF® Dye Antibody Labeling Kit (#92274), and gamma-tubulin primary antibody was conjugated with Alexa 647 fluorophore using Biotium Mix-n-Stain™ CF® Dye Antibody Labeling Kit (#92274). For nuclear staining, Hoescht dye was diluted to 1ug/mL.

### Quantification of planar cell polarity in the node and skin

Celsr1 images were used to generate segmentation masks using Cell Pose software (Stringer et al., 2021) followed by further hand corrections of segmented masks using TissueAnalyzer software for FIJI (Aigouy et al., 2010). Polarity analysis was determined as previously described (Aigouy et al., 2010; Aw et al., 2016). Segmented masks were used in TissueAnalyzer to generate the axis and magnitude of Celsr1 polarity. Rose plots were then generated for Celsr1 polarity using MATLAB.

### Live imaging trachea MCCs

Tracheas were removed from euthanized adult mice. Preparation of trachea for live imaging was performed as previously described (Francis and Lo, 2013). Dissected tracheas were cut in half and placed lumen side down onto a glass bottom dish and covered in ∼100uL of PBS. A square was cut into a circular piece of parafilm and another circular piece of parafilm was adhered to the first piece using nail polish to create a small chamber. This was placed over lumens in the imaging dish. Tracheas were live imaged on a Nikon Eclipse Ti2 equipped with an ORCA-Fusion BT Hamamatsu digital camera.

### Quantification of nodal cilia length and basal body displacement

Nodal cilia were labeled with anti-acetylated α-tubulin and basal bodies were visualized with gamma tubulin. Z-stacks encompassing the full node depth were collected at 1um optical steps. Stacks were processed in ImageJ and analyzed with CiliaQ (Hansen et al., 2021; Schindelin et al., 2012). For each nodal cell the apical boundary (Celsr1) was segmented and the centroid of each node cell recorded. The gamma tubulin position relative to the centroid along the anterior posterior axis was then determined. Basal bodies were classified as anterior, posterior or central/equal. Quantitative data were exported for statistical analysis and plotting in GraphPad Prism 10.

### Zebrafish strain maintenance

The zebrafish *cfap298^tj271^* mutant line, which was previously described (Jaffe et al., 2016), was maintained by outcrossing with wildtype (Burdine lab strain PWT). Transmission of the mutation was determined by PCR and BSTN1 restriction enzyme digest as described previously (Jaffe et al., 2016). To generate *cfap298^tj271^*^/*tj271*^ (*cfap298*^−/−^) homozygous embryos, *cfap298^tj271^*^/+^ fish were crossed. Embryos were maintained at 28° C in E3 (5mM NaCl, 0.17 mM KCl, 0.33 mM CaCl2).

All animal procedures were approved by Princeton’s Institutional Animal Care and Use Committee (IACUC).

### Zebrafish mRNA injection and quantification of phenotypic rescue

mRNA for injections was prepared as previously described (Jaffee et al., 2016). Wildtype *cfap298* mRNA was synthesized from a pSport6.1 vector using a Sp6 mMessage Machine *in vitro* transcription kit (Thermo) while the *cfap298^Δ3aa^* mRNA was synthesized from a pBluescript vector using a T7 mMessage mMachine (Thermo) *in vitro* transcription kit. Embryos from a *cfap298^tj271^*^/+^ heterozygous cross were injected at the 1-cell stage with 500pg of mRNA. Body curvature and heart jogging were scored at 48hpf. Embryos were then lysed in 50mM NaOH boiled at 95°C and genotyped. Embryos were imaged using a Leica M205FA microscope.

---

**Supplemental figure 1.**
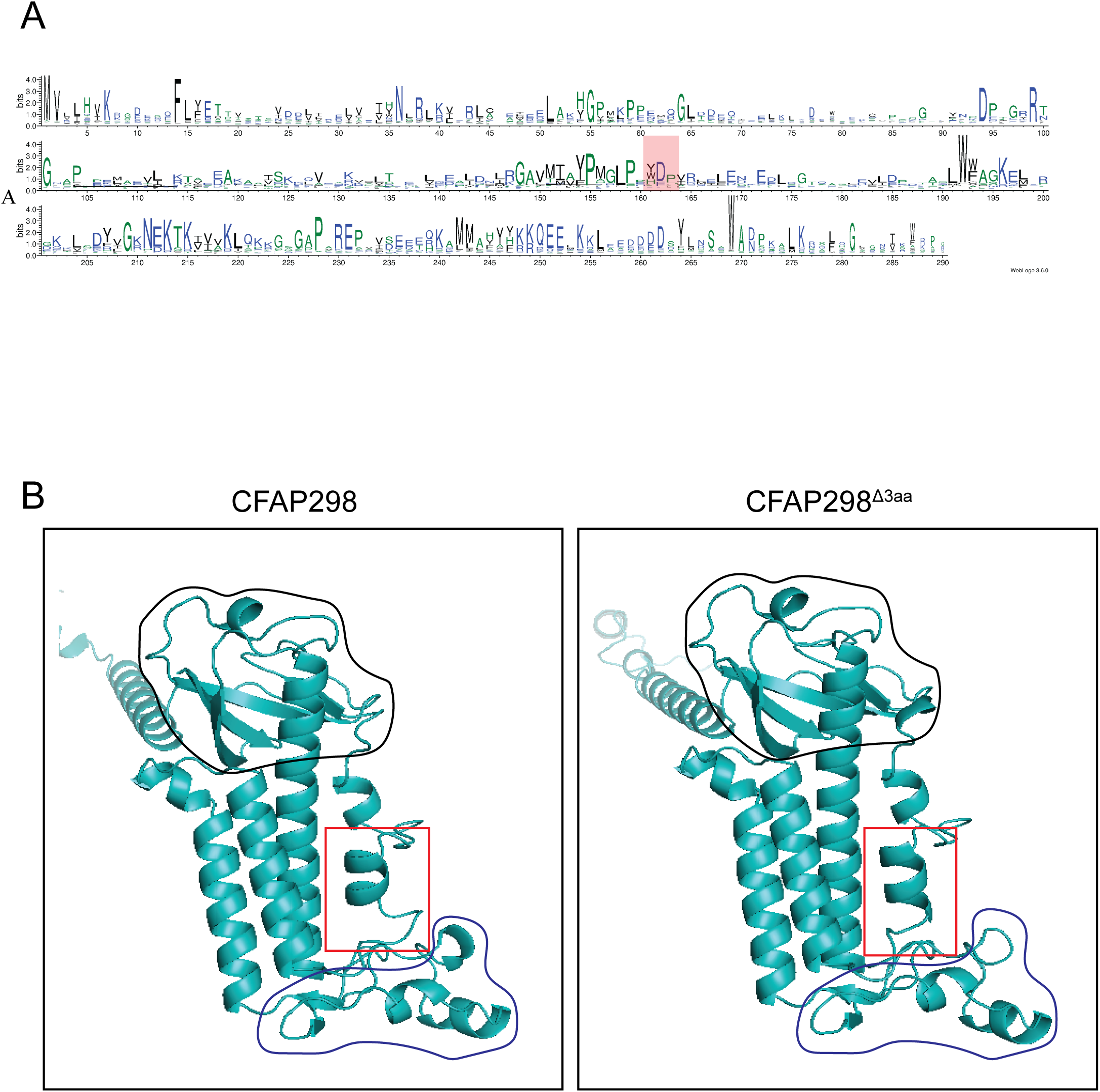
Conservation and AlphaFold predicted structures of Cfap298 and Cfap298^Δ3aa^. A) DMfold multiple sequence alignment logo of Cfap298 from 1610 species. Sequence alignment shows frequency of an amino acid at a given location along the 290 amino acids of Cfap298. Size of the amino acid at a position indicates high incidence of that amino acid at that site suggesting higher conservation. Amino acids 161-163 (orange box) show high conservation across species. B) Alphafold predicted structures of wildtype Cfap298 and Cfap298^Δ3aa^ proteins. Red box indicates the region that includes amino acids 161-163. Black outline indicates the location of the Ubiquitin-like domain. Blue outline indicates location of the Loop domain.

**Supplemental figure 2.**
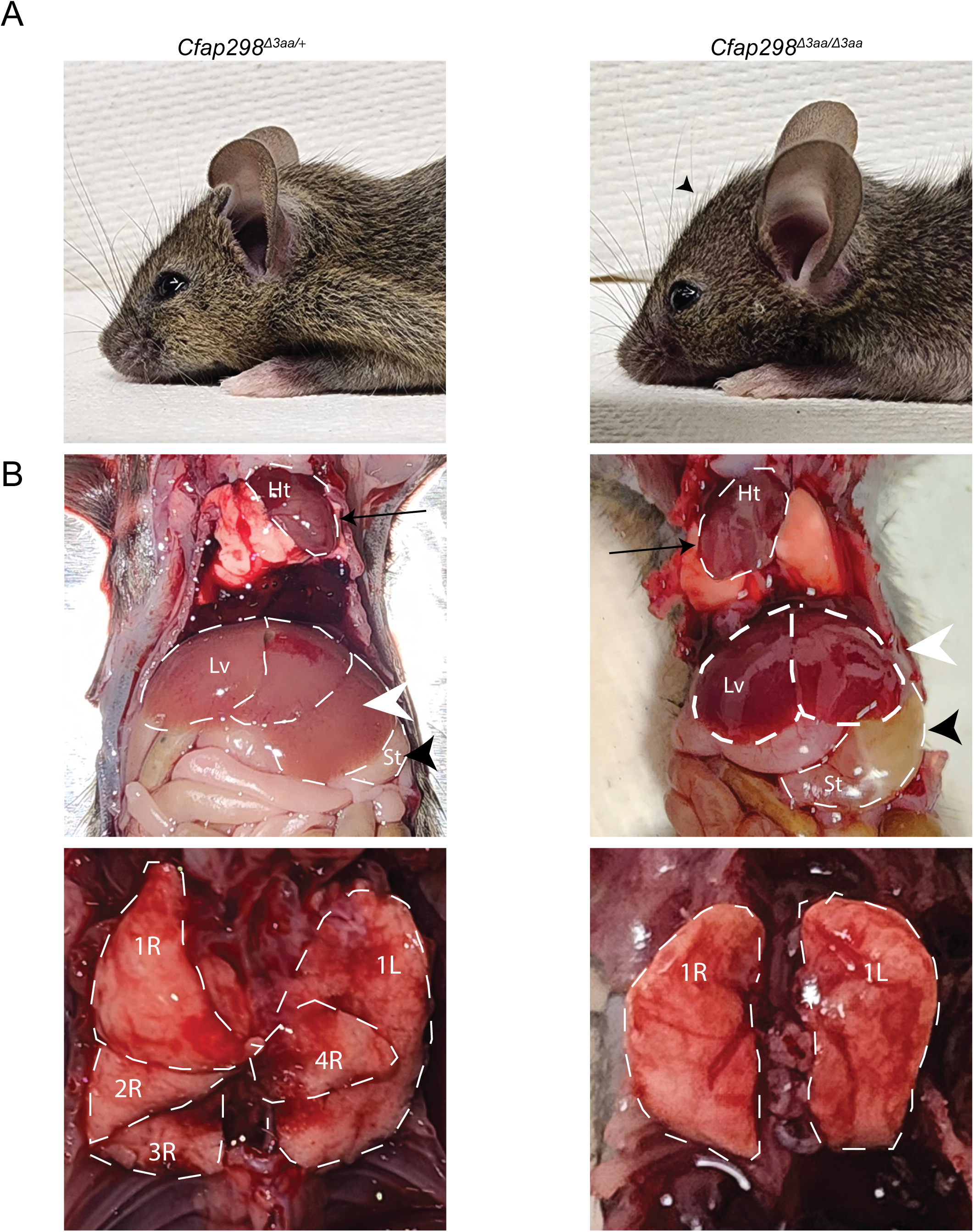
Adult *Cfap298^Δ3aa^* mutant displays cilia motility related defects. A) Side view of *Cfap298^Δ3aa^*heterozygote and homozygous adult mice. Mutant adult shows expansion and doming of the head (arrowhead). B) Representative images of adult necropsies showing left-right defects. *Cfap298^Δ3aa^*^/+^ adult has left-sided heart, three-lobed liver and left sided stomach. Lungs from *Cfap298^Δ3aa^*^/+^ display 4 lobes on the right and 1 on the left. *Cfap298^Δ3aa^* mutant images showing right-sided heart, two-lobed liver, and left-sided stomach. Mutant adult also displays left lung isomerism. Heart (Ht, arrow), liver lobation (Lv, white arrowhead), and stomachs (St, black arrowhead) are outlined. Individual lung lobes are outlined and labeled with their position on either left (L) or right (R).

**Supplemental figure 3.**
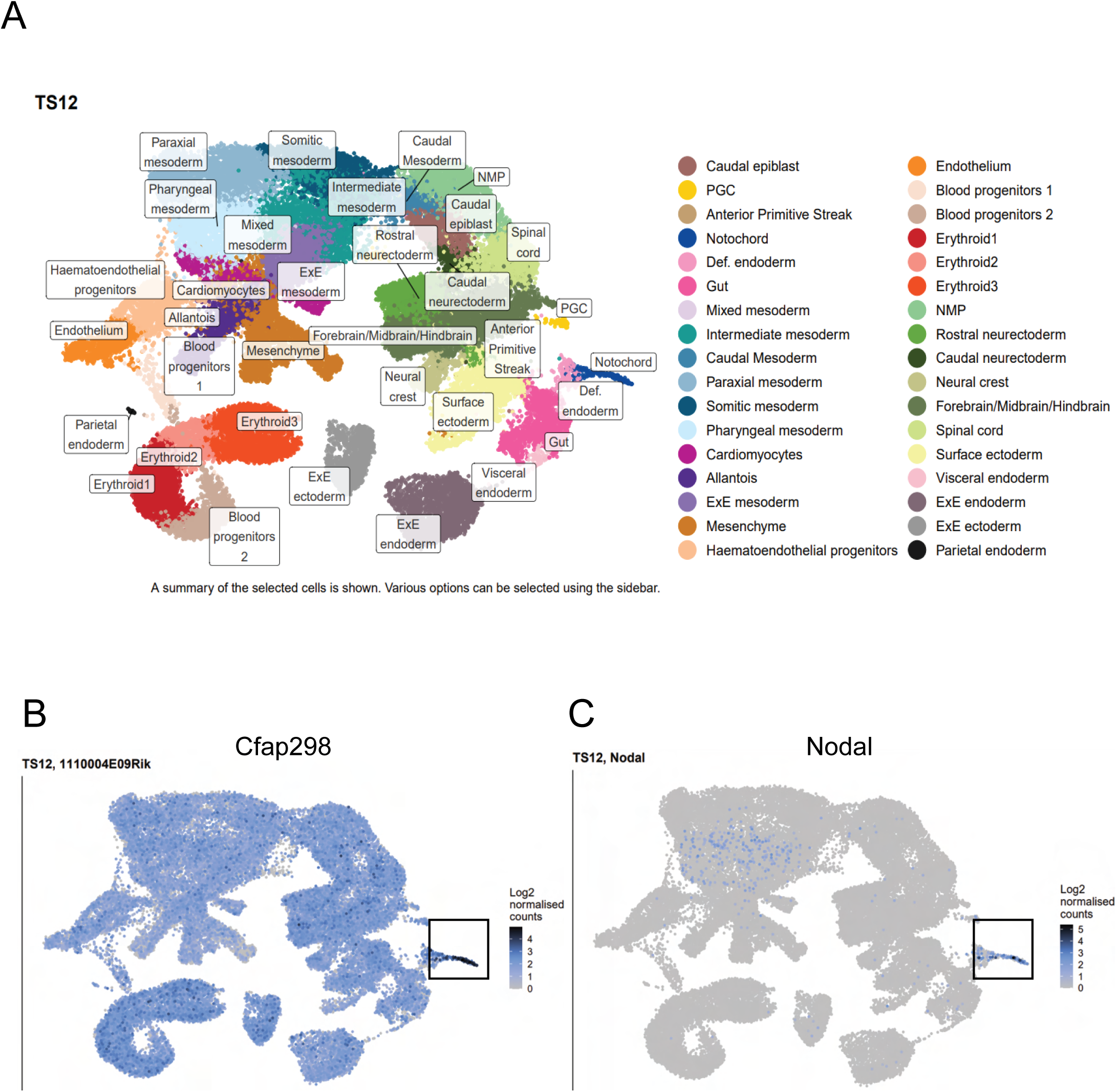
Single-cell RNA sequencing analysis of *Cfap298* expression shows enrichment in the mouse Left-Right Organizer. A) UMAP displaying scRNA seq data from E8.5 embryos (Pijuan-Sala et al., 2019). B) *Cfap298* is expressed in most E8.5 cell clusters and appears to be significantly enriched in the notochord/node cluster (boxed region). C) *Nodal* is specifically expressed within the notochord/node (boxed region) and mesoderm.

**Supplemental figure 4.**
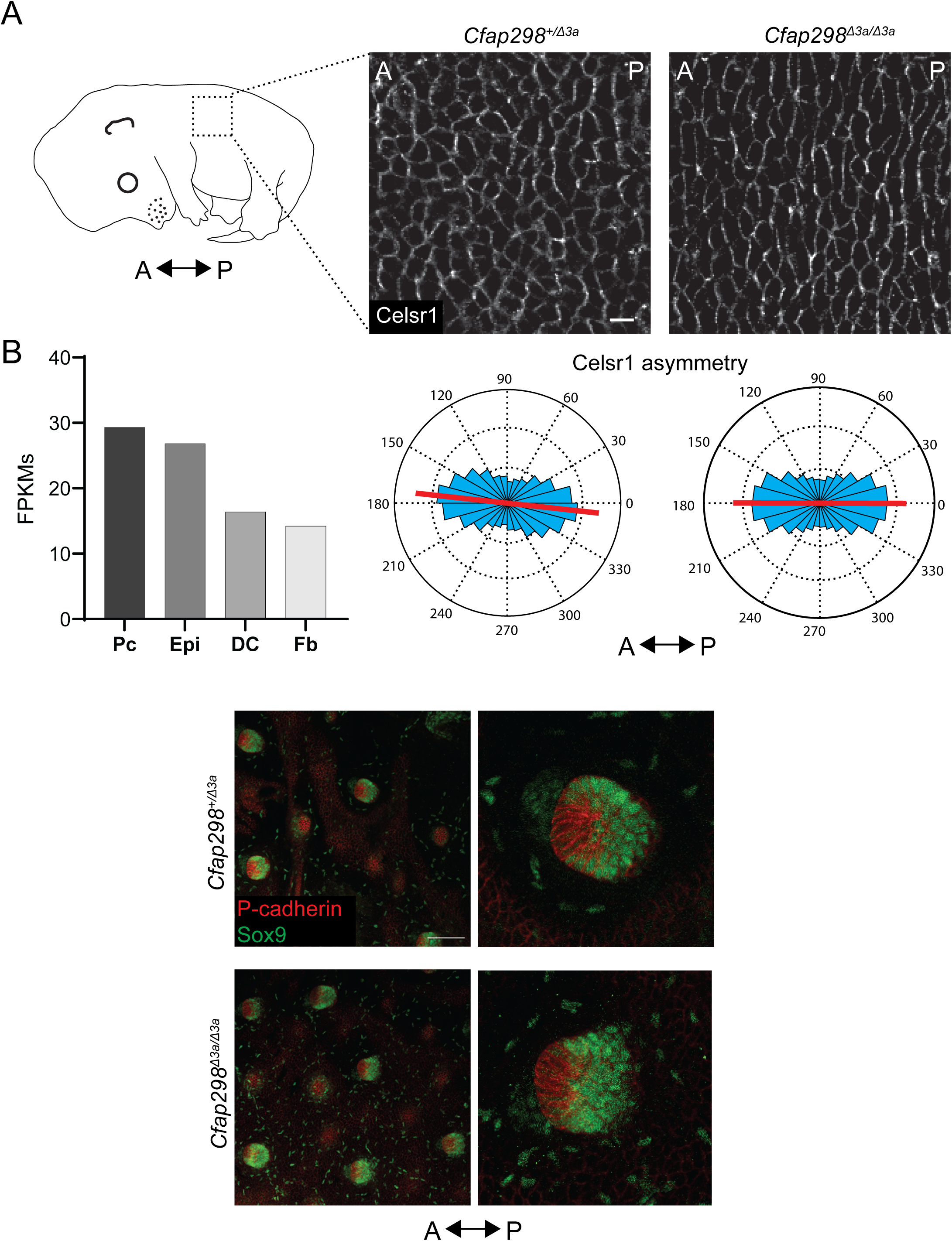
Planar polarity in the mouse epidermis is unaffected in *Cfap298^Δ3aa^*mutants. A) Bulk RNA seq data from E14.5 embryonic skin from (Sennet el al., 2015; Rezza et al., 2016) showing *Cfap298* expression in the placode (Pc), the epidermis (Epi), dermal condensate (DC), and dermal fibroblasts (Fb). Schematic showing region of epidermal skin dissected for Celsr1 staining and quantification in the skin with the anterior and posterior axis (AP) marked along the embryo. Planar view of the basal layer of the epidermis showing Celsr1 staining (grayscale) from E15.5 *Cfap298^Δ3aa^*^/+^ and *Cfap298^Δ3aa^* mutant embryos. Circular histograms depict magnitude and angle of Celsr1 polarity relative to the embryonic anterior-posterior axis for *Cfap298^Δ3aa^*^/+^ (n=3, 11,972 cells total) and *Cfap298^Δ3aa^* mutant (n=3 embryos, 12,313 cells total) embryos. Scale bar= 10um. B) Images of hair follicles from *Cfap298^Δ3aa^*^/+^ and *Cfap298^Δ3aa^* mutant skins with anterior cells marked by P-cadherin (red) and posterior cells marked by Sox9 (green). Scale bar=100um.

**Movie 1: Planar view of MCCs from *Cfap298^Δ3aa/+^* and *Cfap298^Δ3aa^* mutant tracheas.** Videos showing planar views of motile cilia from *Cfap298^Δ3aa/+^* tracheas and immotile cilia in *Cfap298^Δ3aa^* mutant tracheas. Videos were taken at 7fps.

**Movie 2: Side view of MCCs from *Cfap298^Δ3aa/+^* and *Cfap298^Δ3aa^* mutant tracheas.** Videos showing side view of motile cilia from *Cfap298^Δ3aa/+^* tracheas and immotile cilia in *Cfap298^Δ3aa^* mutant tracheas. Videos were taken at 7fps.

